# Context matters: Changes in memory over a period of sleep are driven by encoding context

**DOI:** 10.1101/2022.07.22.501166

**Authors:** Eitan Schechtman, Julia Heilberg, Ken A. Paller

## Abstract

During sleep, recently acquired episodic memories (i.e., autobiographical memories for specific events) are strengthened and transformed, a process termed consolidation. These memories are contextual in nature, with details of specific features interwoven with more general properties such as the time and place of the event. In this study, we hypothesized that the context in which a memory is embedded would guide the process of consolidation during sleep. To test this idea, we employed a spatial memory task and considered changes in memory over a 10-hour period including either sleep or wake. In both conditions, participants (*N* = 62) formed stories that contextually bound four objects together, and then encoded the on-screen spatial position of all objects. Results showed that the changes in memory over the sleep period were correlated among contextually linked objects, whereas no such effect was identified for the wake group. These results suggest that contexts binding different memories together play an active role in consolidation during sleep.

## Introduction

After initial encoding, memories are further processed and strengthened, a process termed memory consolidation. Consolidation occurs during both wake and sleep, with some debate over each state’s unique contribution (e.g., Wamsley and Summer, 2020; Wang et al., 2021). The physiological characteristics of sleep, and specifically non-rapid-eye-movement sleep (NREM), together with the relative paucity of perceptual input that may interfere with processing, are thought to provide an optimal environment for memory consolidation (Diekelmann and Born, 2010; Paller et al., 2021).

Most research on consolidation considered sleep’s role in the evolution of memory for relatively impoverished, isolated stimuli, as is common in memory research. However, real-life memories are rarely isolated, but rather are linked with other memories that were encoded in the same context. Retrieving a specific detail about an event, for example, can produce a plethora of associations and an experience of reliving the full event, a phenomenon termed “mental time travel” (Tulving, 1983). Recollection of a specific detail effortlessly and involuntarily involves the retrieval of other contextually bound details about the same event (e.g., Wheeler and Gabbert, 2017). This memory interrelatedness is fundamental to our understanding of memory in daily living, but little is known about its impact on consolidation in general or on consolidation during sleep in particular.

In this study, we explored whether memories that are contextually bound to one another, and therefore likely to be retrieved together, are also likely to be reactivated together during sleep. The term “context” is notoriously difficult to define, yet most memory researchers agree that it includes spatiotemporal features or other aspects of a remembered event accompanying its defining components (Smith, 1994; Stark et al., 2018; Dulas et al., 2021). Free recall studies that considered the temporal context in which memories were encoded have shown that memories encoded in temporal proximity are more likely to be retrieved together (i.e., the contiguity effect; Kahana, 1996). Retrieval in free recall tasks is also guided by the semantic relatedness between different words, an effect termed semantic clustering (Shuell, 1969; Polyn et al., 2009).

Accordingly, we sought to determine whether contexts driven by temporal or semantic links between memories guide consolidation during sleep as in wake. The experiment contrasted sleep and wake using a between-subject design. Participants used their personal electronic devices at home to create and record unique stories linking different objects with cohesive narratives. Then, they were required to encode the on-screen positions of each object. After a 10-hour delay that either did or did not include nocturnal sleep, they were tested on object positions. We hypothesized that the context in which a memory resided would explain variance in consolidation-related memory changes. Put differently, our prediction was that objects that were linked to the same narrative would have correlated memory trajectories over sleep.

## Methods

### Participants

Participants were recruited from Northwestern University’s academic community, and included paid participants and participants who completed the experiment for course credit. Participants had to have an Android phone and be in the United States while conducting the experiment. In total, 77 participants were recruited (45 men, 31 women, and one genderqueer person; average age = 23.29 years ± 0.53, standard error). Fifteen participants were not included in the final analyses: six participants withdrew before completing the experiment; six participants encountered technical issues; two participants in the Wake group (see below) napped during the day; and one participant completed the final test after more than 12 hours. The final sample included 62 participants (42 men, 20 women; average age = 23.02 ± 0.57 years). These participants were divided into the Wake and Sleep groups (n = 31 each; the Wake group included 20 men and 11 women, average age = 22.97 ± 0.8 years; the Sleep group included 22 men and 9 women, average age = 23.06 ± 0.81 years). All participants consented to participate in the study. The study protocol was approved by the Northwestern University Institutional Review Board.

Participants were randomly assigned to be in either the Wake group or the Sleep group. Both groups underwent the same protocol, with the only difference being the time of day of the two experimental sessions (Figure 1a).

**Figure 1:**
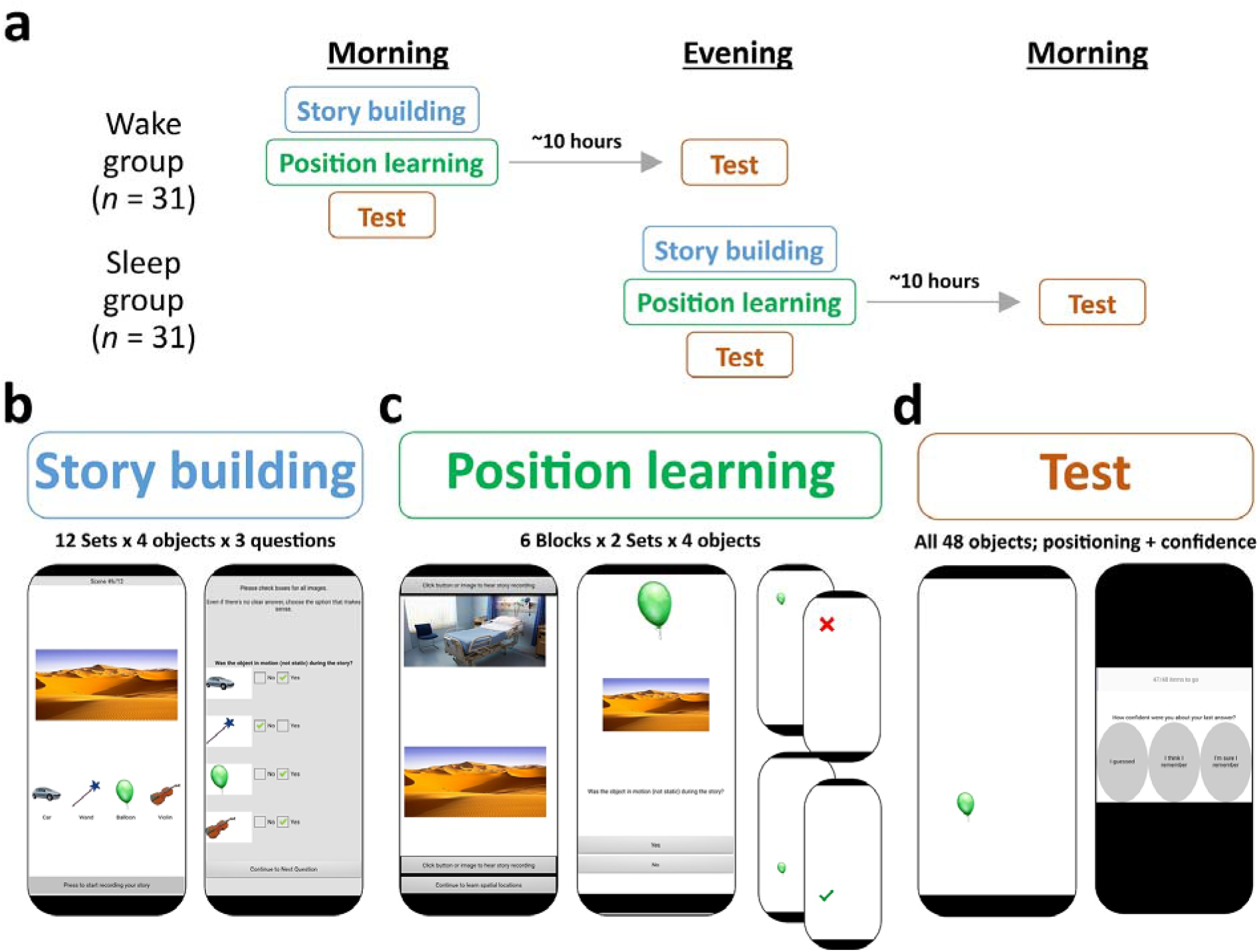
Experimental design. (a) Participants were randomly assigned to either the Wake or Sleep group. (b) In the first session, both groups developed and recorded 12 stories linking a location (e.g., a desert) with four objects. After recording the stories, they responded to three yes/no questions about their stories for each object (the right panel shows one example question). (c) Next, participants engaged in a position learning task. Each object was assigned a random on-screen position. Each block included objects from two contextually bound sets. First, participants were offered a chance to listen to the two stories. After initiating the block, participants were asked in each trial to respond to an object-specific question (middle panel). If they were correct, they attempted to place the object in its correct position. The block continued until all objects were learned to criterion. Feedback was provided in all trials. (d) At the end of the first session, participants were tested on their spatial memory. In each trial, participants also indicated their confidence level. An identical test was conducted in the second session.

### Materials

Participants used their personal Android phones to complete the experiment. A custom application, named “StoryTask,” was designed using MIT App Inventor (Patton et al., 2019). Participants installed the application on their phones and used it to record their audio and touch-screen responses and to present visual and auditory stimuli and instructions. Participants held their phones vertically throughout the task.

Visual stimuli consisted of 48 images of objects and 12 images of places. Object images were square and portrayed either inanimate objects (e.g., a telephone) or animals (e.g., a cat) on a white background. Most images were taken from the BOSS corpus (Brodeur et al., 2010; Brodeur et al., 2014), and some were taken from copyright-free online image databases (e.g., http://www.pixabay.com).

At the core of the experiment was a spatial positioning task, during which participants had to memorize the on-screen positions of images. To standardize the task across devices with different dimensions and resolutions, images were presented within a confined rectangular area of the screen (i.e., the active area). The area was defined as the maximal vertical rectangle that fit within each participant’s screen so that its height will be exactly double its width. The size of the side of each square object image was 20% of the area’s width (i.e., each image occupied 2% of the active area).

Place images portrayed distinct places (e.g., a movie theater; a desert) and were shown horizontally, with a 1:2 proportion between their height and length, respectively. Images were taken from copyright-free online image databases (e.g., http://www.pixabay.com).

Place images were each associated with four objects to create contextually bound sets. Object images were each assigned a random position within the active area. These positions were chosen to be distant from the middle of the screen and any other object’s location (Euclidean distance > 10% of screen width) and were chosen to be at least 10% of the screen’s width from any the active area’s four sides.

### Procedure

Participants were told that the first session would take approximately 90 minutes and the second approximately 20 minutes. They were asked to complete the second session 10 hours after starting the first. Participants in the Wake group were asked to complete the first session in the morning and to avoid napping during the day. Participants in the Sleep group were asked to complete the first session in the evening.

After consenting to participate in the study, participants filled out a set of questionnaires, including the Stanford Sleepiness Scale (Hoddes et al., 1973) and the reduced version of the Morningness-Eveningness Questionnaire (Adan and Almirall, 1991; Loureiro and Garcia-Marques, 2015). Then, they were instructed to download and install the application.

The instructions for the first stage of the task were presented in a video embedded in the application (https://youtu.be/964KR0y7GbU). For this stage (Story building, Figure 1b), participants had to invent a story occurring in the locale depicted in the scene image and involving each of four objects shown. In total, they created 12 stories, each recorded using their device’s microphone. After each story, participants were required to answer three questions for each object: (1) Was the object in motion (not static) during the story? (2) Did the object produce a sound as part of the story? (3) Did the object appear throughout the whole story, start to end? The responses to these questions were conveyed using button presses (Figure 1b, right).

After creating and recording all stories, participants began the second stage of the experiment (Position Learning, Figure 1c). For this task, participants completed six training blocks, each including eight objects that were part of two contextually bound sets. The instructions for this stage were presented in a video embedded in the application (https://youtu.be/ekC1eUnIsC4). Before each block, participants were allowed to listen to the two stories they recorded earlier (Figure 1c, left). Then, they were shown each object in its assigned on-screen position. Next, they underwent a continuous, multi-trial learning task to encode each object’s position. Each positioning trial began with a presentation of the object image along with its associated location (e.g., balloon, desert; Figure 1c, center) and one of the three questions presented previously. The participant had to answer that question correctly (i.e., as indicated during the story-building stage) to continue to the next part of the trial, and had 7 seconds to respond by pressing “yes” or “no.” In the next part, participants attempted to recall each object’s on-screen position within a 7-second response interval. Recall was deemed correct if the position indicated by the participant was within a short distance of the true position (less than 20% of the active area’s width). As feedback, the object appeared in the true position. The next trial then ensued. Each block consisted of repeated loops of trials with the drop-out method. Objects were considered learned if they were correctly positioned in two consecutive trials, and learned objects were dropped from the following loop. A block ended when this learning criterion was achieved for all objects.

After learning, participants had to take a break for at least 5 minutes before starting the next stage (Test, Figure 1d). Here, participants tried to place each object in its true position. Objects were presented in a pseudorandom order and no feedback was provided. In each trial, participants had 7 seconds to position the object. After each trial, participants indicated their confidence level on a 3-level Likert scale (“I guessed,” “I think I remember,” “I’m sure I remember”). After positioning all 48 objects, participants were tested on object-location associations. For each object, four images of locations were presented, including the location previously presented with the object. Participants attempted to indicate which location was linked with each object. This test concluded the first session.

The application was designed so that participants would be unable to start the second session until at least 6 hours have passed since completing the first session. In the second session, participants first filled out another questionnaire, and then began a test that was identical to that of the first session (including the object-scene association test). After completing the second session, participants were instructed to email their data to the experimenter, erase the data from their device, and uninstall the application.

### Statistical analyses

Data were analyzed using Matlab 2018b (MathWorks Inc, Natick, MA). Intraclass correlations with missing values were calculated using the irrNA (version 0.2.2) package in R (version 4.1.2).

To account for differences in screen sizes, the sizes of all visual stimuli were proportional to the participant’s screen size and spatial accuracy was estimated using units normalized to the screen size. Memory performance was assessed by fitting mixed linear models. Memory for individual objects was considered in these analyses, accounting for random intercept effects for different participants. An ANOVA was used to report the statistical significance of the components of the model, and dummy variables were used for comparisons between conditions. Table 1 includes the models used in this analysis. Some analyses were conducted on a subset of objects based on the ordinal confidence levels (e.g., limited to the “guessed” trials). In these cases, only objects rated with those confidence levels on both the pre-sleep and post-sleep tests were considered.

**Table 1:**
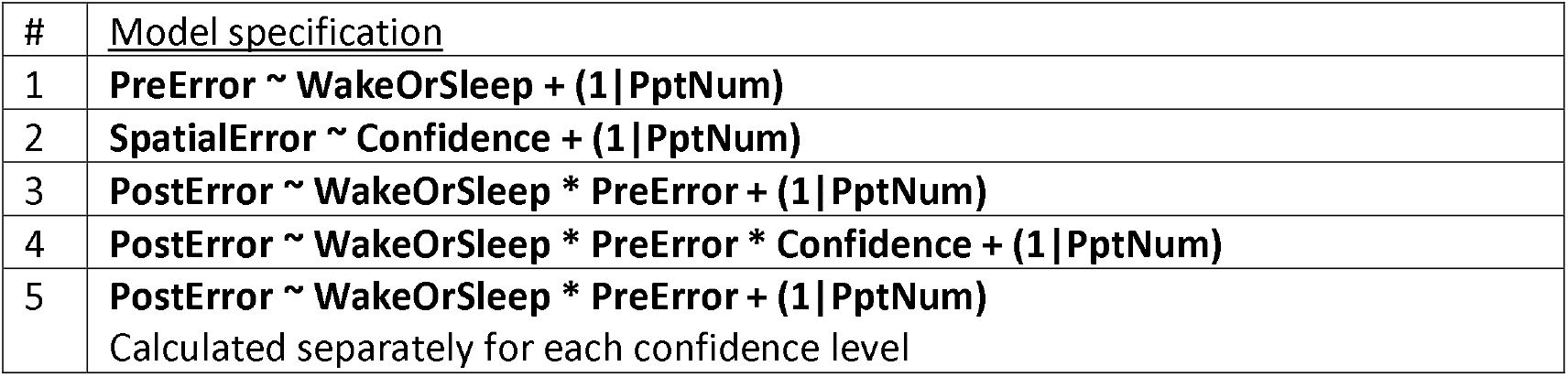
Mixed linear models used in analyses. SpatialError – spatial error in a test; PreError – spatial error in the first experimental session; PostError – spatial error in the second experimental session; WakeOrSleep – categorical group indicator; PptNum – categorical participant indicator; Confidence – ordinal confidence level.

Our main hypothesis was that variability in memory trajectories would be explained by shared contexts. To test this hypothesis, we used intra-class correlation (Koo and Li, 2016). This metric, ICC, is symmetrical (i.e., whereas *inter*-class correlations predict Y from X, *intra*-class correlations predict how clustered together different values of X are) and can be used to calculate the correlation between more than two values. We used the (1, k) form of ICC (Shrout and Fleiss, 1979; Koo and Li, 2016). For object positions that were not rated by participants as guessed, we calculated the change in positioning error over the delay. We then calculated the ICCs for each participant to consider two sub-hypotheses: (1) to test whether semantic clustering explained the variability in the changes in memory over the delay period, we considered ICC for objects linked within the same contextually bound set; (2) to test whether temporal context explains the variability in the changes in memory over the delay period, we calculated the mean change for each contextually bound set (i.e., four objects) and then used an ICC analysis to test whether those are correlated within block (i.e., whether performance for two sets linked within the same training block were correlated). The ICCs obtained through these analyses were compared with the ICC results obtained through permutation tests with mixed labels (n = 10,000) for each participant. We conducted a one-tailed paired t-test to test whether the true ICC was larger than the average ICC value calculated using the permutation test. In addition, we used a one-tailed two-sample t-test to test whether the true ICC for the Sleep group was higher than that of the Wake group. Analyses that did not include object-level measures of performance were conducted using two-tailed two-sample t-tests.

## Results

Participants were randomly assigned to Wake and Sleep groups (*n*=31 each; Figure 1a). The groups followed the same protocol, which included engaging in two experimental sessions, with the second session starting approximately 10 hours after the first. The Wake group trained in the morning and were then tested in the evening, whereas the Sleep group trained in the evening and were tested in the morning. Training consisted of a story building stage (Figure 1b) and a position learning stage (Figure 1c). In the story building stage, participants encoded contextually bound sets, which included an image of a location linked with four images of objects. In the position learning stage, they learned the on-screen positions of the objects. Learning was organized into six blocks, each including objects from two contextually bound sets which were learned in temporal proximity. Participants were tested on object positions twice – once shortly after learning and once after the delay period (Figure 1d).

The Wake and Sleep groups did not differ in terms of age [*t*(60) = 0.08, *p* = 0.93], Morningness-Eveningness scores [*t*(60) = 1.47, *p* = 0.15], or the length of the delay between the first and second sessions [*t*(60) = 0.33, *p* = 0.74]. The Stanford Sleepiness Scale assessed before the beginning of the first session showed higher sleepiness for the Sleep group relative to the Wake group (2.29 vs 3.48, respectively; *t*(60) = 4.29, *p* < 0.001). To consider whether differences in fatigue or time of day (i.e., circadian effects) might have impacted learning or memory performance on the first session, we compared positioning error rates for the first session’s test between groups and found no significant differences [*F*(1, 2815) = 1.06, *p* = 0.30; Sleep group = 15.42% ± 1.3, Wake group = 17.27% ± 1.3; Model #1 in Table 1].

### Memories recalled at intermediate confidence levels benefited more from sleep than wake

In their tests of spatial recall, participants were required to indicate their confidence level in each trial (Figure 2a). As expected, error rates were lower as confidence levels increased [*F*(2, 5713) = 445.16, *p* < 0.001; Guess = 26.04% ± 0.9, Think = 17.32% ± 0.8, Know = 10.68% ± 0.8; Model #2 in Table 1, Figure 2b]. To test whether sleep improved memory in this task, we used a model to predict memory on the second session based on pre-delay error rates and group (Wake vs Sleep; Model #3 in Table 1). In this analysis, a main effect of group would indicate a uniform effect of sleep/wake, and an interaction between pre-delay errors and sleep would indicate that the effect of sleep/wake depended on the initial strength of the memory. Our results indicated that neither effect was significant [*F*(1,2757) = 2.2, *p* = 0.14 for the main effect of group; *F*(1, 2757) = 0.2, *p* = 0.66 for the interaction].

**Figure 2:**
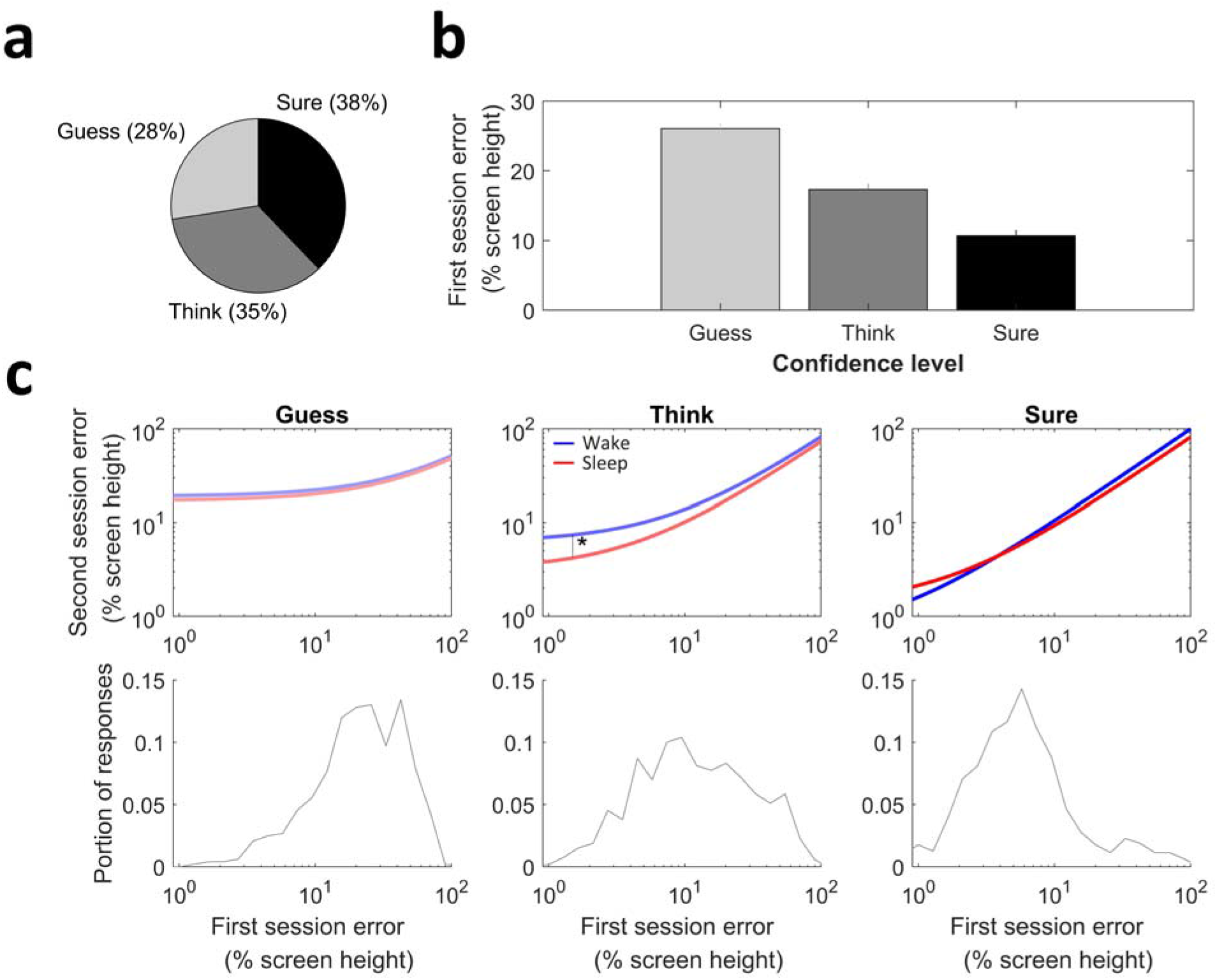
Memories recalled at moderate confidence levels benefited from sleep. (a) Distribution of confidence as rated by participants. (b) Average error rates for each confidence level. Error bars represent standard errors of the mean for all objects. (c) The effects of sleep on memory for objects rated with different confidence levels. Each of the upper panels shows the error rates for the first and second sessions on the X and Y axes, respectively (log-log scale). The lines show the linear correlation between first and second session errors (note that lines seem curved due to the log-log axes). For objects with intermediate confidence level, the sleep group showed significantly lower post-sleep errors. The bottom panels show histograms of the first session errors. * - *p* < 0.05.

In an exploratory analysis, we next incorporated confidence levels into the analysis to test whether the effect of sleep on memory for object positions interacts with confidence levels. We therefore used a model to predict memory on the second session based on three factors: memory on the first session, confidence levels, and group (Wake vs Sleep; Model #4 in Table 1). As expected, both memory on the first session and confidence levels, as well as this interaction, were positively correlated with memory on the second session (all *p* values < 0.001). Interestingly, two significant interactions suggested that confidence levels drove memory benefits: the interaction between group and confidence level [*F*(2, 2749) = 6.65, *p* < 0.01]; and the interaction between group, confidence level, and memory on the first session [*F*(2, 2749) = 3.5, *p* < 0.05]. The effect of group and the interaction between group and memory on the first session were not significant (*p* > 0.26).

To resolve the interactions, we conducted analyses separately for each confidence level (Model #5 in Table 1; Figure 2c). All three models found that memory on the first session significantly predicted memory on the second session (all *p* values < 0.001). However, only the objects rated with the “think” confidence level showed a significant effect of sleep, indicating overall greater memory benefits of sleep relative to wake [*F*(1,525) = 6.18, *p* < 0.05; Figure 2c, center]. In addition, these objects also showed a trend toward an interaction between group and memory on the first session, indicating a possible differential effect of sleep on memory for objects based on their initial memory strength [*F*(1,525) = 3.05, *p* = 0.08]. In other words, results indicated that sleep improved memory for intermediate confidence objects, with a trend toward greater improvement selectively for objects with good pre-sleep accuracy. No significant effects emerged for the objects rated with the “guess” confidence level (all *p*-values > 0.38) or the “know” confidence level (all *p*-values > 0.12).

### Variability in memory benefits over sleep is explained by shared context

To investigate the role of context in the consolidation of memories, we considered the change in memory over the delay between the first and second sessions (i.e., the memory trajectories). Our analytic approach was to leverage the variability in trajectories to evaluate the impact of shared contexts. If the context binding memories together plays some active role during the delay period, we expected contexts to explain some of the variability in trajectories. More specifically, we hypothesized that context would drive consolidation during sleep. Therefore, we hypothesized that memory trajectories for objects linked within the same contextually bound sets (i.e., interlinked within the same story) would be more correlated than chance if that delay included sleep (Figure 3a). We did not have an *a-priori* hypothesis regarding the impact of a wake delay of similar duration, but if sleep has a privileged role in memory consolidation, then trajectories would be less correlated after wake relative to sleep.

**Figure 3:**
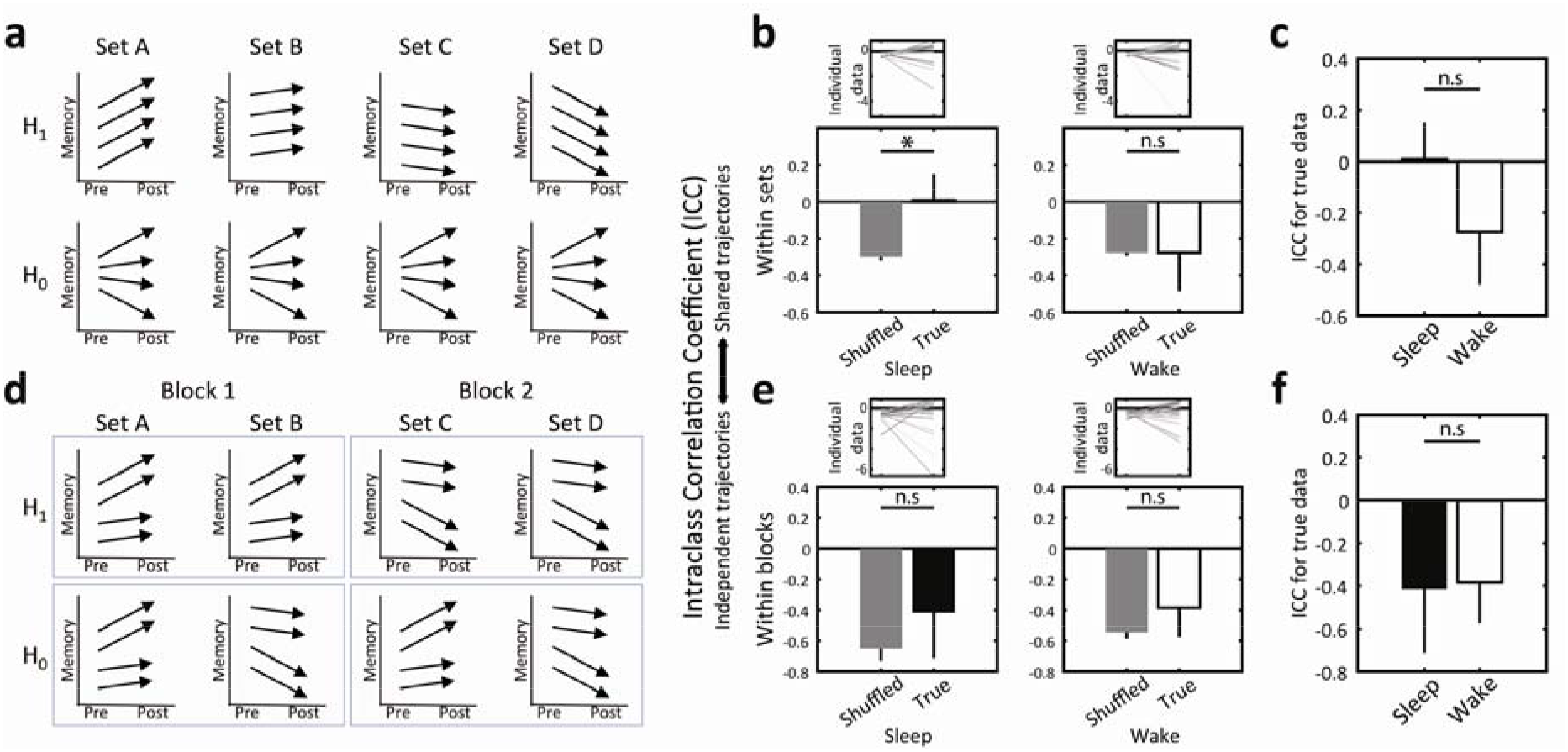
Variability in memory benefits over sleep is explained by shared semantic context. (a) Analytic approach. We hypothesized that binds between objects linked within the same contextually bound sets would drive changes in memory performance over sleep. If this were the case, memory trajectories (i.e., changes in memory between the first and second session; signified by arrows) would be correlated within sets (as shown in H_1_) for the sleep group. (b) Intraclass correlation coefficients were calculated to estimate within-set correlations per participant. These values were compared with the average correlations obtained through permutation tests. The upper panel shows individual participant values for the Sleep and Wake groups (left, right respectively). The bottom panels show the across-participant averages. (c) Direct comparison between the correlation coefficients for the Sleep and Wake groups. (d) We hypothesized that the temporal context binding together contextually bound sets that were learned within the same blocks would drive changes in memory performance over sleep. If this were the case, average memory trajectories within sets would be correlated within blocks (as shown in H_1_) for the sleep group. (e) Intraclass correlation analyses to consider the effect of temporal context on memory. Designations follow those introduced in panel b. (f) Direct comparison between the correlation coefficients for the Sleep and Wake groups. Error bars signify standard errors of the mean across participants in all panels. * - *p* < 0.05; n.s - *p* > 0.05.

To test this hypothesis, we considered all objects that were not designated as “guesses” in our analysis. For each participant, we calculated the intraclass correlation coefficient, a measure of overall agreement between different values within a group. This measure, ICC, reflects how clustered together contextually bound memory trajectories are. We compared these “true” values to the average ICC values obtained by shuffling the labels in 10,000 different permutations for both the Sleep and Wake groups (Figure 3b). Our results showed that the Sleep group indeed showed higher-than-chance ICCs [*t*(30) = 2.11, *p* < 0.05]. The Wake group did not show a similar effect [*t*(30) = 0.01, *p* = 0.5]. Finally, we compared the “true” ICCs for the Sleep and Wake group and found no significant difference between the two [*t*(60) = 1.13, *p* = 0.13; Figure 3c]. Taken together, these results suggest that memories that share a semantic context are consolidated together during sleep.

To explore whether a similar effect can be observed for temporal context (i.e., with the temporal proximity between memories at encoding driving consolidation benefits), we leveraged the structure of our task. Each block during the position learning stage included two contextually bound sets which were learned within temporal proximity of one another (Figure 1c). We therefore hypothesized that memory trajectories for objects within one set would be correlated with the trajectories of the set learned within the same block in the Sleep group (Figure 3d). Like before, we did not have an *a-priori* hypothesis regarding the Wake group, except that context would have a lesser impact on delay-related changes on that group relative to the Sleep group.

The analytic approach employed to test this hypothesis was similar to the one used to test within-set intraclass correlations. The average memory trajectories were calculated per set and then submitted to an ICC test to consider within-block correlations for each participant. These results were compared to those obtained using a permutation test. Unlike for semantic contexts, our results did not support our hypotheses. Both in the Sleep group and in the Wake group, true ICC values were not significantly different than those obtained in the permutation test [*t*(30) = 0.78, *p* = 0.22; *t*(30) = 0.76, *p* = 0.23, respectively; Figure 3e]. Additionally, ICC values were not significantly higher for the Sleep versus the Wake group [*t*(60) = −0.09, *p* = 0.54; Figure 3f]. Taken together, our results did not support the hypothesis that temporal context plays a role in consolidation during sleep.

## Discussion

In this study, we investigated whether the encoding contexts of memories impact the manner in which they are consolidated over a 10-hour delay. Objects bound together by unique encoding contexts were tested before and after a delay that either did or did not include nocturnal sleep. Results showed that sleep improved retrieval only for memories rated with an intermediate level of confidence. Our analyses considered two different types of contexts – semantic contexts (i.e., memories that shared meaningful narrative connections with one another) and temporal contexts (i.e., memories that were encoded within the same time interval). We found that some of the variability in the changes in memory over the delay were explained by semantic context only if the delay included sleep. Conversely, we found that temporal context did not significantly explain the variance in consolidation benefits in wake nor sleep.

These results complement previously reported findings from our group demonstrating that manipulating consolidation using external cues during sleep impacts contextually bound memories (Schechtman et al., 2022). Whereas that study utilized methods to bias reactivation selectively towards certain memories in a nap setting, the current study did not involve a causal manipulation, instead focusing on the consequences of nocturnal sleep involving spontaneous, endogenous memory reactivation. In addition, this study included a wake control that allowed us to probe the specific interaction between context and sleep. Encouragingly, the two studies together converge on the conclusion that context guides memory processing during sleep. Moreover, a central limitation of the current study – that it reveals changes in correlation patterns but falls short of demonstrating causality – is overcome by the other study from our group. Likewise, a central limitation of the study of Schechtman et al. (2022)—that it involves cued rather than spontaneous reactivation and may therefore not reflect the cognitive benefits of non-manipulated sleep—is overcome by the present study.

Our results, showing a benefit of sleep only for memories rated with an intermediate level of confidence (“think” vs “guess”/”know”), diverge from previous findings exploring the relationship between memory strength and consolidation. Previous studies have revealed that sleep is especially beneficial for weakly encoded memories (e.g., Drosopoulos et al., 2007; Diekelmann et al., 2010). If this were the case in our study, we would expect the greatest sleep benefits for object locations recalled with the lowest confidence. A general difficulty in considering the question of memory strength across multiple experiments is that differences between tasks and cognitive demands make comparisons extremely challenging. It could be, for example, that memories in the intermediate confidence zone in our study would have been rated as weakly encoded in the context of another study. Are “weakly encoded” memories defined in a relative way (i.e., the weakest memories for a given task) or in an absolute way (i.e., based on some task-independent metric, such as exposure time or depth of processing)? This question has not been thoroughly investigated. Finally, it is worth mentioning that it has been hypothesized in the past that perhaps sleep preferentially benefits memory in the intermediate range (Stickgold, 2009, Figure 4), as in our study.

As with many studies comparing sleep with wake, our study has several notable limitations. First, our design does not allow us to disentangle the beneficial effects of sleep from the detrimental effects of wake interference. The changes over a delay period involving sleep may have nothing to do with sleep itself, except for it being a period of time that is less cognitively demanding and prone to interference relative to a similar period of time spent awake. Second, the circadian differences between the two groups (i.e., the time of day of the first and second session) may have contributed to the differences between them. Although we have tried to rule this explanation out by analyzing the effects of time of day on performance, this factor may still have had some contribution to the observed results. Finally, our null results with regard to the effects of temporal context on consolidation should be interpreted cautiously. Despite the present findings, the idea that temporal encoding factors influence consolidation should not be ruled out. Our design intentionally emphasized semantic context in its task demands, whereas temporal contexts were encoded incidentally. The structure of our experimental blocks may have also hampered the operationalization of temporal context by adding many strong event boundaries within blocks (e.g., breaks between trials). More research should be conducted to address the role of temporal context on consolidation during sleep.

Our results suggest that memories are not consolidated independently of one another during sleep, but rather that the context in which they were encoded and the relationships between memories play a role in the consolidation process. As research studies in cognitive neuroscience increasingly include more naturalistic designs, there should be a growing emphasis on incorporating more of the complexity of memory interrelationships along with richer environments. We see our results as another step towards acknowledging that memory processes cannot be understood devoid of their overarching contexts – during both wake and sleep.

## Acknowledgments

This work was supported by NIH grant K99-MH122663 and by NSF grants BCS-1921678 and BCS-2048681.

## Author contribution

All authors contributed to the design of this study and helped revise the manuscript. E.S and J.H collected the data. E.S conducted the analyses and wrote the initial draft of the manuscript.

